# Landscape structured by physical settings and benthic polychaete and avifauna habitat uses in a mangrove-vegetated estuary

**DOI:** 10.1101/2019.12.12.874008

**Authors:** Shang-Shu Shih, Tzung-Su Ding, Chang-Po Chen, Shou-Chung Huang, Hwey-Lian Hsieh

## Abstract

Mangrove expansion monopolizes estuarine landscapes by diminishing habitat diversity and hence biodiversity. Physical landcover types, including mangrove vegetation, influence polychaete and avifauna habitat uses. The connections between the physical to biota-associated landscapes warrant investigation. We determine how to best describe the landscape in a mangrove-vegetated wetland according to the physical, polychaete and bird domains and identify what physical attributes would affect the biota-associated landscapes. Differences among the physical and biota-associated landscapes were evaluated using multivariate ordination analyses. Six physical landcover types were aligned along elevation, inundation and sedimentary gradients. The polychaete-associated landscape was structured by three landcover types, mainly mangroves and tidal flats with intermediate and high inundation. Deposit-feeding spionid and nereid, carnivorous goniadid and suspension-feeding sabellid polychaetes depended on the different landcover types. Shorebirds occurred distinctively in tidal flats with large, open surface areas. Egrets characterized tidal flats and mangroves, and foliage and ground gleaners characterized mangroves. Open tidal flats are crucial to polychaetes, which are the main prey of shorebirds and are also important to egret foraging. Our results suggest that effective management strategies for conserving these migratory birds require the maintenance of open tidal flats in the landscape.

## Introduction

Essential components of a healthy mangrove ecosystem include mud and sand flats, tidal waterways, shallow water areas and circulating waters in addition to mangrove stands [1, 2]. These mosaic and interconnected landcover types make mangrove ecosystems varied in terms of landscape function and the production of both terrestrial and aquatic organisms [3-5]. Furthermore, mangrove ecosystems are also among the most threatened ecosystems on Earth due primarily to the devastating effects of anthropogenic activities [6, 7] and natural disturbance [8]. Assessments of the relationships between the supplies provided by mangroves and demand from human society have demonstrated intimate bottom-up and top-down connections between the functions of mangrove ecosystems and the services they offer to human wellbeing [9]. When considering mangrove trees alone, they compose a simple ecosystem with limited vegetation niches and low bird species richness [10, 11]. At the landscape level, however, landscape heterogeneity within both mangrove ecosystems and their surroundings is crucial in characterizing mangrove-dependent bird assemblages [11-13]. Mangroves have been found to have both positive and negative roles in ecosystems. For example, as they act as both foundation species and ecosystem engineering species, mangroves can substantially change landscape structure through their biohydrological attributes. These changes could promote habitat availability for some species but not for others [14]. To properly manage mangrove-dominated wetlands, the data necessary for understanding the interactions among the geomorphological, hydrological, biological and socioeconomic domains that underlie mosaic landscapes are still missing for many of these wetlands, and a lack of relevant knowledge has caused many mangrove rehabilitation efforts to fail [1].

Most suggestions regarding mangrove-associated landscape management have focused on avifauna and the anthropogenic effects of different types of land use on bird communities [11-13]. There are few studies reporting the landscape- and/or physical habitat-based connections between avifauna and their food sources [15]. Macrobenthos polychaetes and bivalves and fishes, for instance, are the main preys of shorebirds and egrets [15-17], while insects are a major food source of foliage gleaners [18]. Polychaete and bivalve distributions are largely controlled by local physical driving forces, including hydrology, geomorphology and sedimentology [19-23], while insect distributions are tightly associated with vegetation composition [24]. These findings suggest that bird distribution must closely follow the benthos and vegetation structure. Consequently, the context of a local landscape arises as a result of the physical setting and macroinvertebrate- and bird-specific landscapes. Information on these interconnected landscapes and human land uses would greatly improve conservation efforts focusing on mangrove-vegetated wetlands.

The Wazihwei Nature Reserve is located in the Danshuei River estuary, which hosts the northernmost population of the mangrove *Kandelia obovata* in Taiwan [25, 26]. This reserve was designated in 1994 to preserve the mangrove trees, which are strictly protected by Taiwan’s Culture Heritage Reservation Act. This area is also an important wintering and stopover site for migratory shorebirds, egrets, and other waterbirds [27]. From 1984 to 2013 (30 years), some shorebird and egret populations remarkably declined by several tens to hundreds of times, while during approximately the same time period, the mangrove-vegetated area increased by approximately 30% [27]. These changes suggest that the decreasing waterbird abundance in this reserve might be attributed to the simplified landscape structure as a result of mangrove overexpansion [2].

To resolve the conflicts between mangrove protection and biodiversity enhancement, it is necessary to understand whether identifiable connections between the physical and biotic landscapes exist and what physical attributes affect such connections. Using the Wazihwei wetland as a case study site, the purposes of the present study were to assess (1) how the best landcover types within the physical landscape and biotic landscape are characterized, in other words, how well the landscape is described by the physical and biotic domains, and (2) what physical attributes contribute to structuring the biota-coupled landscape. These assessments will promote our understanding and sound management of mangrove-vegetated wetlands from a landscape perspective.

## Materials and methods

### Study area

The study area is approximately 33.3 hectares in size and is located in the Wazihwei Nature Reserve adjacent to the mouth of the Danshuei River estuary in northern Taiwan (Fig 1, [27]). In the estuary, the M2 tide is the primary tidal constituent, with a mean tidal range of 2.3 m and up to 3.3 m during spring tides. From west to east, there were four main landcover types: mangrove vegetation, intertidal mud and sand flat, tidal creek and sand dune. The mangrove vegetation covered approximately 11.2 hectares and was composed of a single species, *Kandelia obovata*. The form of the tidal creek changed from its original meander into almost a straight line along with the expanding mangroves [28]. Anthropogenic activities included fishing boat anchoring, sightseeing on the walking trail and wastewater discharge from the upland residential communities and industries [27]. The construction of cement-paved roads and walking trails in the sand dune area had affected the local hydrodynamics and sediment transport, consequently shifting the outlet of the tidal creek to the south and forming a curved sand spit to the southwest [28]. The plants found in the sand dune area consisted primarily of wormwood (*Artemisia capillaris*), beggar’s tick (*Bidens pilosa* var. *radiata*), beach morning glory (*Ipomoea pescaprae* subsp. *brasiliensis*) and sea hibiscus (*Hibiscus tiliaceus*) [29].

**Fig 1.**
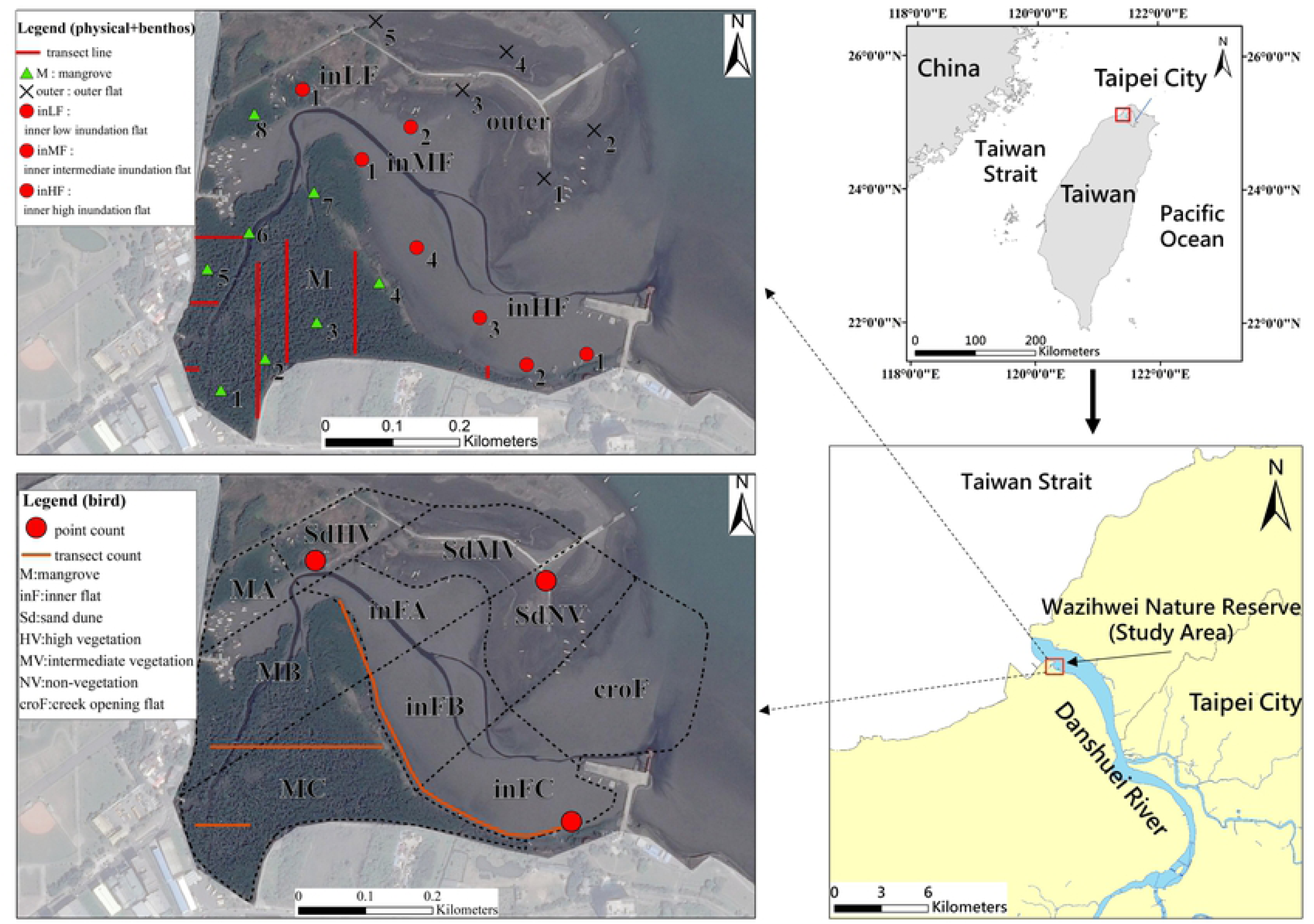
Sampling stations established in the studied Wazihwei Nature Reserve for the collection of physical attribute and polychaete and avian assemblage data. Seven line transects were deployed under the mangrove canopy for water level measurement using a communication-vessel system. Dashed lines represent the 10 survey zones established for collecting bird variable data.

### Sampling schemes for physical attributes and biotic assemblages

A grid composed of 6 × 8 transects was deployed, and sampling stations were established where these transects intersected (Fig 1). The distance between stations was approximately 100 to 165 m. Data on physical variables and the polychaete assemblage were collected simultaneously from all stations. Eight stations were established in the mangrove-vegetated zone. The inner intertidal flat was subdivided into areas that experience low (inLF), intermediate (inMF) and high (inHF) inundation according to the previously observed inundation frequency (Chen CP and Shih SS, personal observation), and 1 to 4 sampling stations were established within each of these areas. Five stations were established in the outer intertidal flat. Data on the physical variables at each of these 20 stations were collected every 3 months from October 2013 to July 2014. Thereafter, from January 2015 to October 2015, the number of sampling stations was reduced to 9: 2 in inHF, 1 in inMF, 3 in the outer intertidal flat, and 3 in the mangrove zone. As a result, a total of 116 samples (20 stations × 4 times + 9 stations × 4 times) were collected. The same sampling scheme was used to sample the polychaetes, but the sampling period was only from October 2013 to July 2014, resulting in a total of 80 samples.

The avifauna of the entire study area was surveyed along 5 × 4 transect lines that were roughly perpendicular and parallel to the river channel (dashed lines in Fig 1). These lines delineated three landcover types (mangrove, intertidal flat and sand dune). The numbers of areas surveyed were 3 in the mangrove vegetation (MA, MB, MC), 3 on the inner flat (inFA, inFB, inFC), and 1 in the outlet of the primary tidal creek on the flat (croF). Based on vegetation coverage (Huang SC, personal observation), the sand dune zone was subdivided into areas with high, intermediate and no vegetation (SdHV, SdMV, SdNV, respectively). The avifauna surveys were conducted monthly from October 2013 to November 2015 (except from October to December 2014) and included a total of 230 surveys (10 areas × 23 times).

### Measurements of physical attributes

The geomorphological and hydrological variables measured included the exposed open surface area of the intertidal flat when the tides had completely retreated, elevation, slope, inundation frequency, flow resistance, and flow velocity. The topography of the nonmangrove area was investigated using a TOPCON Total Station (GTS226). The adjacent Tenth River Management Office, Water Resources Agency, Ministry of Economic Affairs, Taiwan, was used as the benchmark reference. The benchmark elevation and water stage records were obtained from the Taiwanese fundamental benchmark of Keelung. This fundamental benchmark was adopted as the zero orthometric height of Taiwan. Furthermore, a communicating-vessel system (CVS) was established under the mangrove canopy [27]. The water levels of the CVS were used to measure exact heights above the substratum surface across the mangrove stand area. The area where the topography significantly varied required more measurement points along an established transect. Seven transects were established in the CVS survey (Fig 1). The elevation data from the mangrove and nonmangrove areas were then input into ArcGIS to produce topographic contour maps. Data indicating the slope and exposed open intertidal surface areas were obtained from this contour map and water stage records. A HOBO Water Level Logger (model U20-001-01) was installed in a nonmangrove location to monitor the water stage every 30 minutes. The Weibull method was employed to analyze the inundation frequency associated with the different water levels [30].

To acquire the flow characteristics, flow velocity and water depth of the reserve to evaluate different flow fields, a horizontal two-dimensional hydrodynamic model, RMA2, was used [31]. Flow resistance, including friction drag and pressure drag, was calculated by multiplying the drag coefficient, object projected area (herein mangrove trees) and square of flow velocity [32]. The annual daily flow discharge of the Danshuei River was set as the upstream boundary of the model, while the hourly water level recorded at the river-mouth gauge station during ebbing (from high tide to low tide) was considered as the downstream boundary. The Manning’s n values of the mangrove and nonmangrove areas were set as 0.08 and 0.03 [33, 34]. The eddy viscosity was set to 20 m^2^ s^-1^ considering the shallow and slow flow in wetlands [35].

The measured sediment variables included grain size, silt and clay content, sorting coefficient, moisture content and pH. Sediment cores were collected from each station using an acrylic tube with a diameter of 2.6 cm. During transport to the laboratory, all sediment samples were kept cool at approximately 4°C. Granulometry was determined following a protocol developed by Hsieh and Chang (1991) [36]. In the laboratory, interstitial water was obtained after the sediments were centrifuged, and its pH values were measured using a pH meter (Mettler Toledo InLab 437). Moisture content was calculated as the percent weight loss after the sediments were oven dried at 60°C to a constant weight. Expressions of units for all physical attributes are given in S1 Table.

### Measurement of biotic assemblages

Polychaete assemblages were sampled using a PVC corer with a diameter of 10 cm that was pushed approximately 10 cm deep into the sediment. Then, the contained sediment was sieved through a 0.5 mm screen. The specimens retained on the screen were relaxed in menthol and fixed in 90% ethanol. The polychaete specimens were identified to the lowest taxonomic level, and the numbers of individuals in each polychaete taxon were counted. Polychaete densities were expressed as the number of individuals m^-2^.

The avifauna was surveyed using transect and total count methods. In the mangrove-vegetated zone, transect counts, which heavily depend on the detection of bird sounds, were performed along the transects at a walking speed of 1-1.5 km hr^-1^. Total counts were conducted in open areas where birds could be directly observed. Three total count locations, two at approximately the northern and southern tips of the inner tidal flat and one in the sand dune area, were established (Fig 1). Locations of bird individuals were recorded with the aid of 8 × 25 binoculars and a 20 × 60 telescope. The bird surveys were conducted for approximately 1 to 1.5 hrs during ebb tides at daytime. The recorded bird species were divided into 6 guilds: shorebirds, egrets, waterfowl, ground gleaners, foliage gleaners and aerial predators (listed in S2 Table).

### Statistical analyses

To improve normality for the multivariate analyses, we transformed the physical attribute and biotic variable data. The transformation formulae are listed in S1 Table. Only species in the polychaete assemblage constituting more than 2% of the total abundance or showing greater than 2.5% occurrence in all samples were included in the multivariate analyses to reduce the influence of rare species on the ordination [37]. A total of 6 polychaete species (see S6 Table) and 6 avian guilds were used in the analyses. The included polychaetes commonly occur in the Danshuei River estuary [37].

We used the multivariate ordination technique of canonical discriminant analysis (CDA, [38]) to distinguish the best landcover types on the basis of the physical attribute and biotic assemblage variables. This analysis was also used to identify which variables constitute the principal components differentiating the landcover types. The abundances of polychaete species and the abundance and species richness of bird guilds with correlation coefficients with the Can1 and Can2 variables greater than 0.40 were considered important components and included in subsequent analyses.

We also used other ordination methods, canonical correspondence analysis (CCA) and redundancy analysis (RDA, [38]), to examine the relationships among the biotic variables and physical attributes. In analyses of the relationships for the bird assemblages, only the datasets that simultaneously contained both bird variables and sedimentary attributes were used. In addition, because each bird survey zone (Fig 1) was large and included several stations used for collecting sedimentary attributes, the values of the sedimentary attributes for each bird survey zone were the means of those attributes collected from the given stations in that given zone. A prior principal component analysis (PCA) or detrended correspondence analysis (DCA) was separately conducted with the data for the polychaete or avian assemblages to assess the gradient lengths of the first PCA or DCA axis. The lengths of the gradients were 4.9 and 1.6 (in SD units) for the density of the polychaetes and the abundance plus species richness of the bird assemblages, respectively. Therefore, the unimodal model (CCA) was appropriate for the polychaete assemblages, while a linear model (RDA) was more appropriate for the bird assemblages [39]. We used the automatic forward-selection mode in our analyses and only included environmental or so-called explanatory variables that explained a significant proportion of the remaining variation based on a Monte Carlo test with 999 permutations at p < 0.1. The CDA was performed using SAS software 9.4 [40], while the ordination analyses (CCA and RDA) were conducted using CANOCO for Windows v.5.0 [41].

## Results

### Physical attributes

On average, the exposed open intertidal surface area was 18,517 m^2^; the elevation was 0.74 m above the mean sea water level; the slope was mild and with 0.06 inclination; the inundation frequency was 24%; and the flow velocity was slow, at 0.06 m sec^-1^, while the flow resistance was 0.12 N m^-2^ (S3 Table).

The sediments consisted of poorly sorted fine sand with moderate amounts of silt-clay and moisture while the interstitial water was approximately neutral. The grain size averaged 130 μm, the silt-clay content was 38.7%, the sorting coefficient was 1.63, the moisture content was 30.6%, and the pH was 7.06 (S3 Table).

### Abundance and species richness of the biotic assemblages

Thirteen polychaete species, including one unknown species among the juvenile nereids, were recorded, with a mean density of 440.7 individuals m^-2^ (Table 1). The capitellid *Capitella* sp., the spionid *Malacoceros indicus* and the nereid *Neanthes glandicincta* were among the most abundant species, with densities of 147.4, 96.2, and 80.1 individuals m^-2^, respectively. These three species represented 73.5% of the whole polychaete assemblage. The remaining polychaete species exhibited lower densities, ranging from 1.6 to 36.8 individuals m^-2^.

**Table 1.**
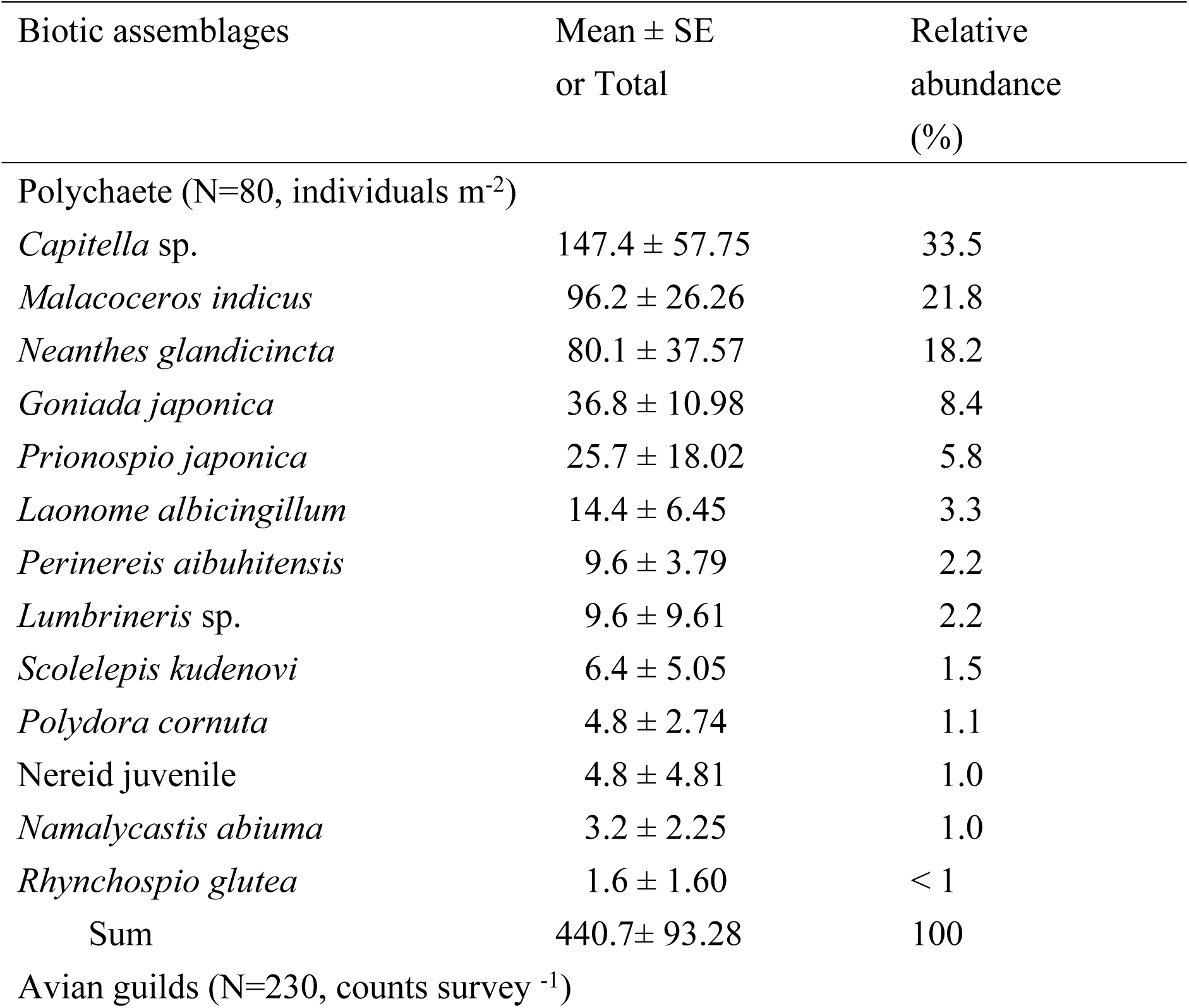
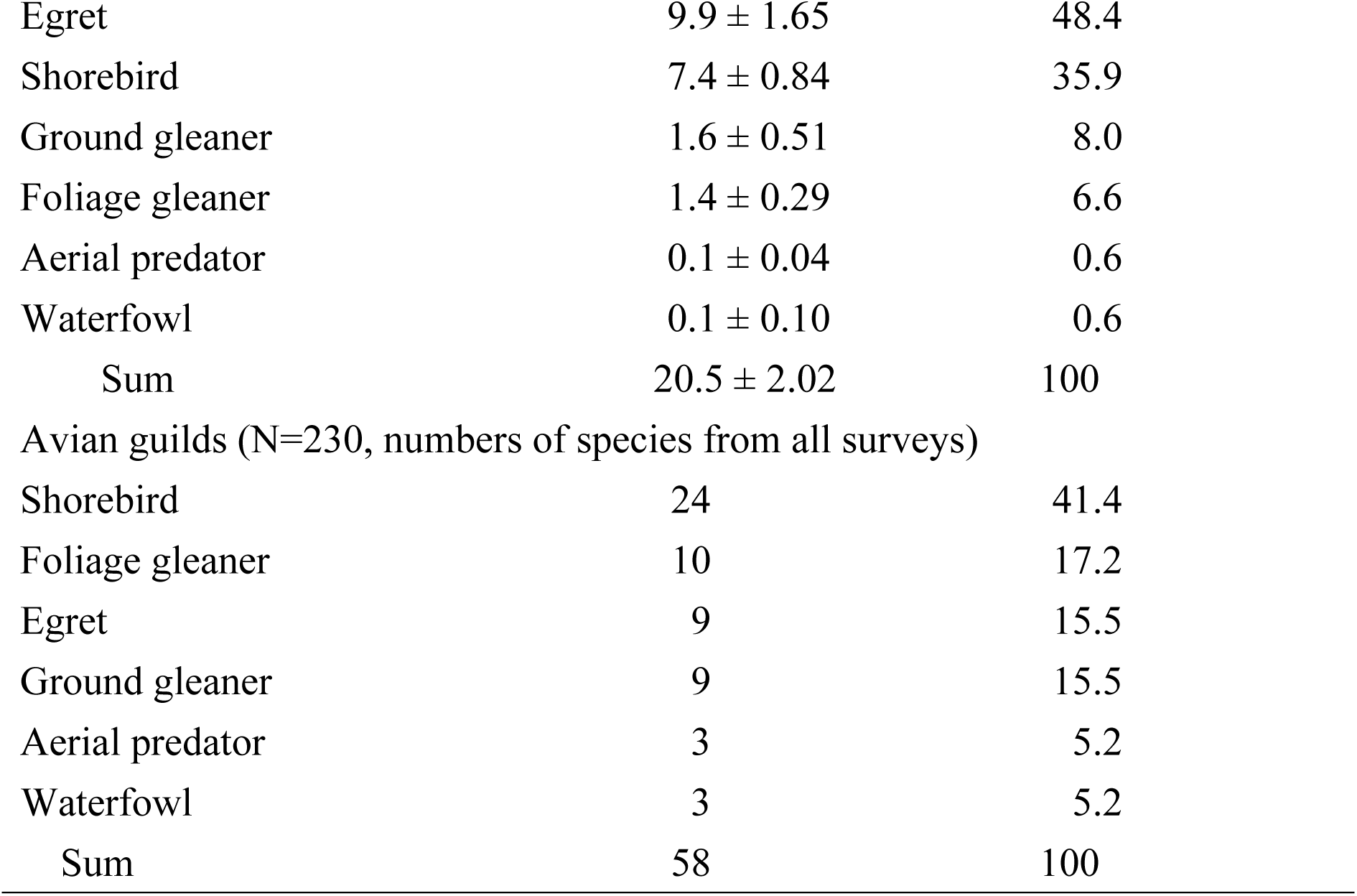
Mean and relative abundance of polychaete and avian assemblages and avian species richness in the Wazihwei Nature Reserve wetland in the Danshuei estuary, northern Taiwan during 2013-2015

A total of 4715 bird individuals and 58 bird species were recorded (S2 Table). Egrets and shorebirds were the two most abundant guilds, with an average of 9.9 and 7.4 counts per survey, respectively. These two guilds combined accounted for 84.3% of the total bird counts (overall average: 20.5 counts, Table 1). The ground and foliage gleaners presented fewer counts, and the aerial predators and waterfowl were even rarer. Twenty-four species of shorebirds were recorded and constituted 41.4% of the total bird species (Table 1).

### Landcover types distinguished according to physical attributes

According to the CDA based on the physical attributes, six landcover types were well distinguished (Table 2). These landcover types were the outer flat (outer), mangroves, inner low inundation flat (inLF), inner intermediate inundation flat (inMF), and inner high inundation flat (inHF) (Fig 2A, S4 Table). The latter was further separated into two subtypes: one included Stations 1, 2, and 3, and the other included Station 4. Station 4 was located at a much higher elevation and had greater flow velocity and resistance than Stations 1, 2, and 3 (S5 Table).

**Table 2.** Summary of the results of the canonical discriminant analysis (CDA) performed on physical attributes and polychaete and avian assemblages

| Variables | Wilks'<br>lambda<br>p | Cumulative<br>% variance<br>explained<sup>a</sup> | Canonical correlations |  |  |  |  |  |  |  |
| --- | --- | --- | --- | --- | --- | --- | --- | --- | --- | --- |
|  |  |  | r<sub>1</sub> | Test |  |  | r<sub>2</sub> | Test |  |  |
|  |  |  |  | Approximate<br>F | df | p |  | Approximate<br>F | df | p |
| Physical attributes | < 0.0001 | 96 | 0.95 | 32.66 | 32, 385 | < 0.0001 | 0.92 | 19.22 | 21, 302 | < 0.0001 |
| Polychaete | < 0.0001 | 84 | 0.80 | 8.06 | 24, 245 | < 0.0001 | 0.68 | 5.85 | 15, 196 | < 0.0001 |
| Avifauna Abundance | < 0.0001 | 97 | 0.72 | 8.60 | 30, 878 | < 0.0001 | 0.48 | 3.54 | 20, 730 | < 0.0001 |
| Species richness | < 0.0001 | 95 | 0.78 | 12.53 | 30, 878 | < 0.0001 | 0.56 | 5.55 | 20, 730 | < 0.0001 |

280 <sup>a</sup>: from combination of Can1 and Can2

**Fig 2.**
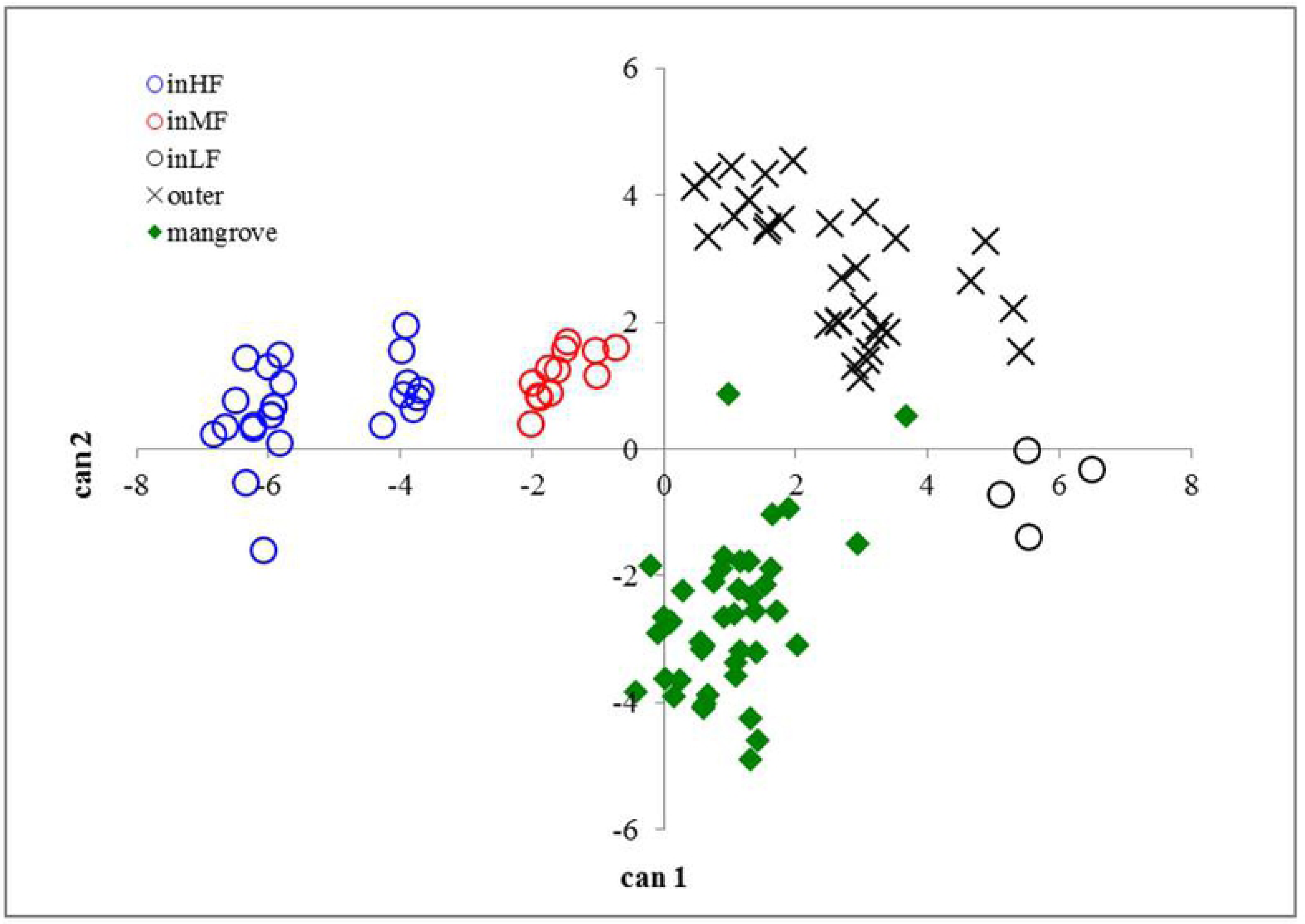
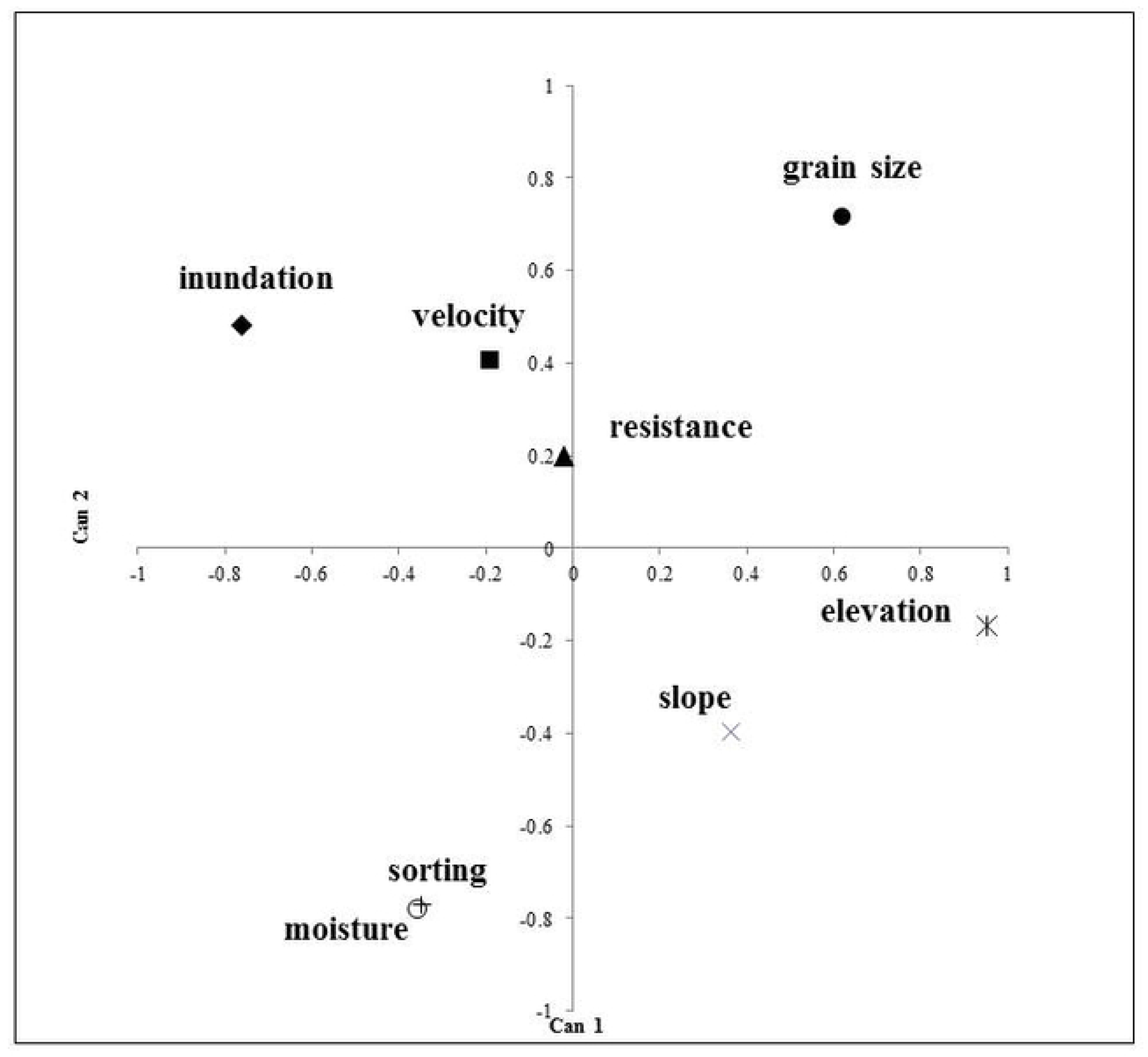
The results of canonical discriminant analysis for physical landcover types. (A) Differentiation of six physical landcover types. (B) Ordination of physical attributes. Landcover type abbreviations as in Fig 1. Note that for the inner high inundation flat (inHF), the small cluster at the right of the larger cluster consists of samples from Station 4.

In terms of the differentiation of landcover types, on Can1, elevation and inundation frequency were the most important variables, and these two factors were inversely correlated. On Can2, the sorting coefficient, moisture content and grain size were the most important attributes, and the latter variable was inversely correlated with the former two attributes (Fig 2B). These ordinations showed the existence of two intercorrelated zonal gradients across the landscape. One was decreasing elevation with increasing inundation, and the other was decreasing grain size with decreasing sorting degree but increasing moisture content (Fig 2). The outer flat, inner low inundation flat and mangroves were located at higher elevations in the northern, northeastern and southwestern regions relative to the lower-elevation inner intermediate and high inundation flats in the central southern regions (0.95 to 1.97 m above vs. 0.04 m below mean sea level). The inundation became prolonged across these regions in a trend parallel to the inclination in which the inner high inundation flat was covered with water more than 5-fold longer than the inner low inundation flat and mangrove vegetation (54% vs. none and 10% of the time, respectively).

In terms of the sedimentary gradient, the sediment in the outer flat and inner low inundation flat consisted of sand and was moderately to poorly sorted (0.84 to 1.56), with larger grain particles (191 to 272 μm) and less moisture (approximately 19%) in the northern and northeastern regions than in the southwestern and southern regions (Fig 2B, S4 Table). In addition, the sediment in the mangrove and inner intermediate and high inundation flat areas was composed of mud (44 to 67 μm) with high moisture (approximately 40%) and was poorly sorted (1.75 to 2.08).

### Association between landcover type and polychaetes

According to the CDA, the density distributions of polychaete species were associated with three distinguishable landcover types (Table 2). These landcover types were primarily the inner high and intermediate inundation flats as well as the mangrove vegetation (Fig 3A). In addition to these three separable landcover types, the inner intermediate inundation flat also occurred in combination with mangrove vegetation in several locations, while mangrove vegetation was also associated with all of the outer and inner low inundation flats and some locations of the inner intermediate inundation flat.

**Fig 3.**
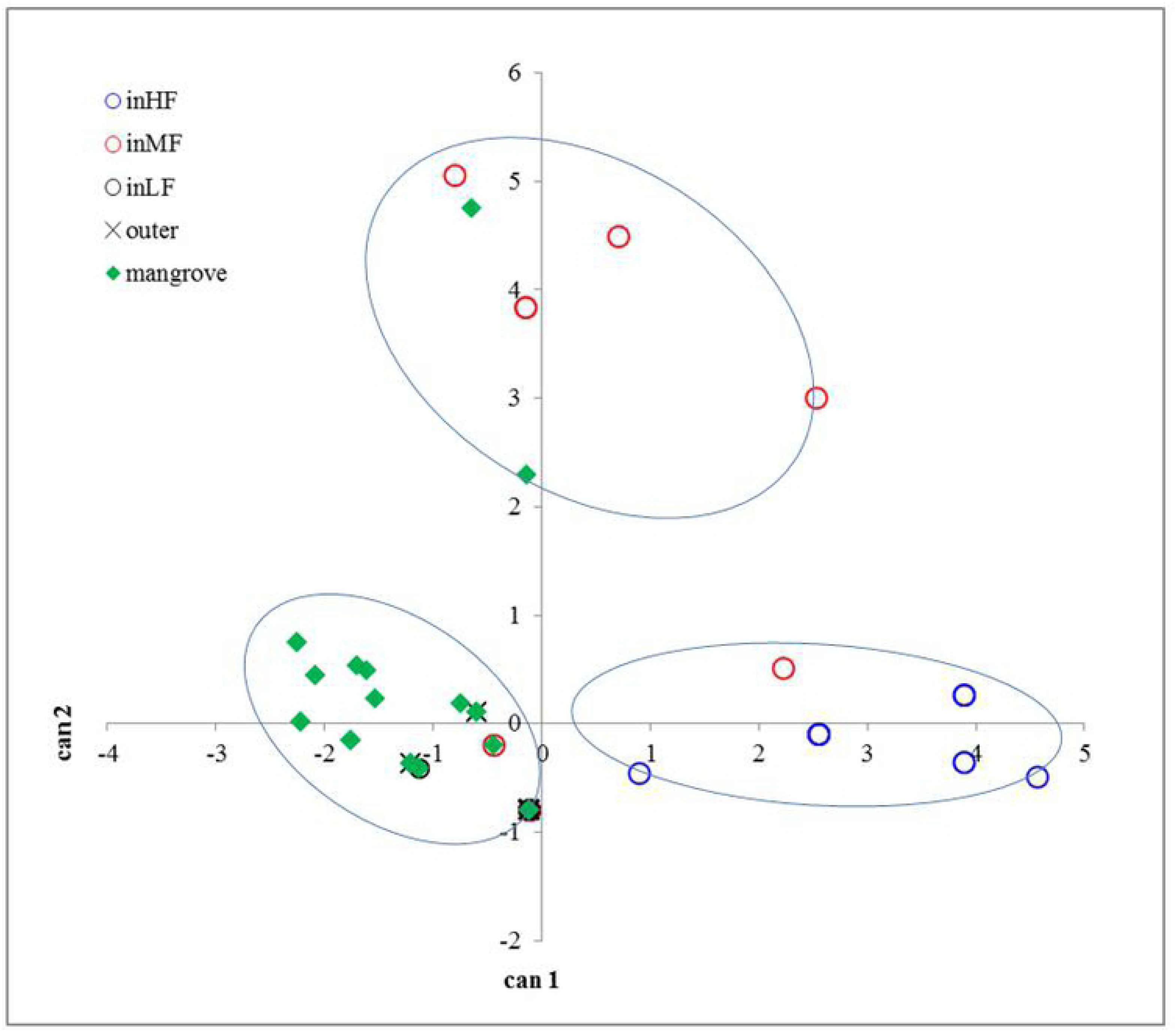
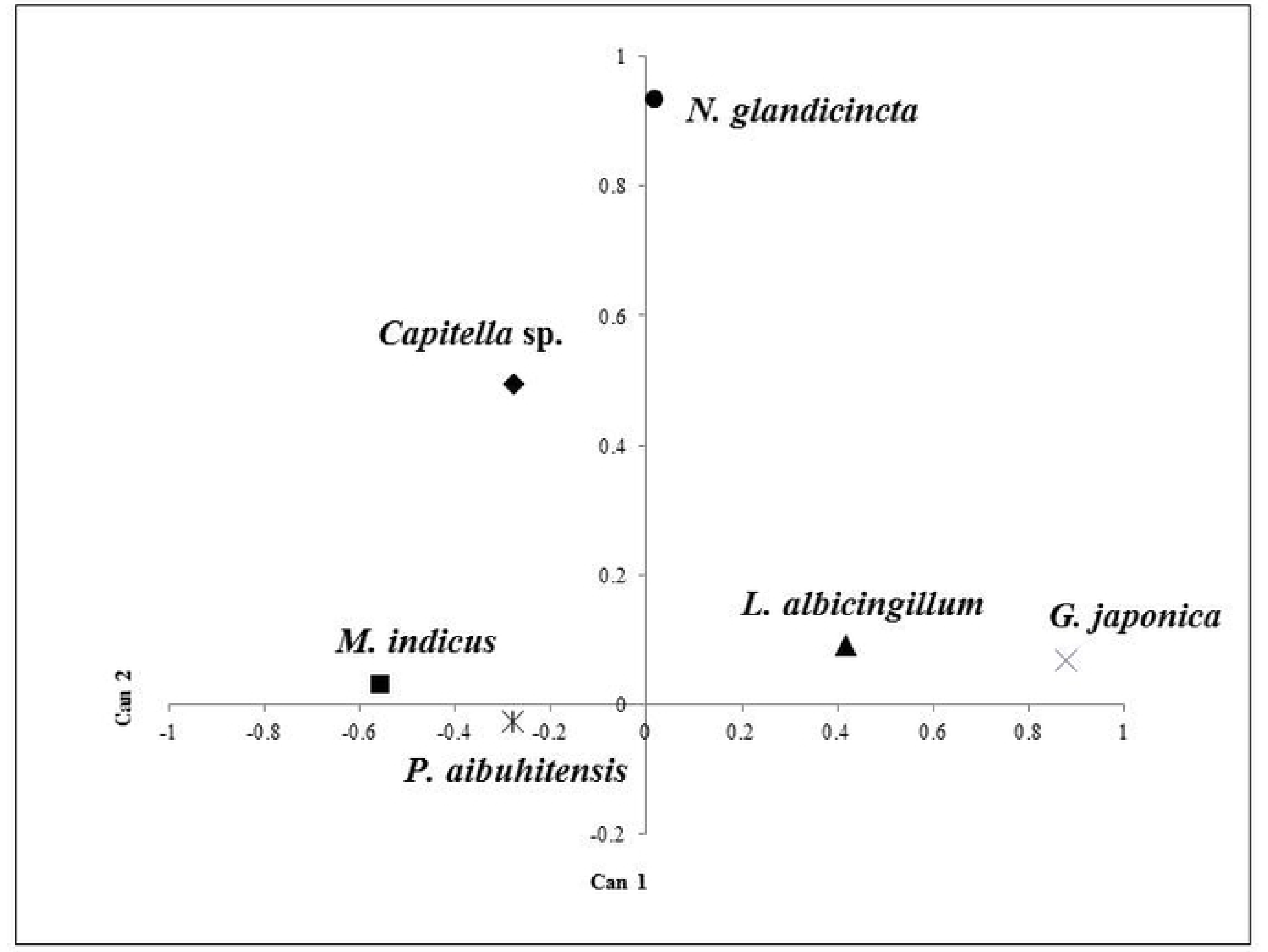
The results of canonical discriminant analysis for polychaete assemblage. (A) Differentiation of three polychaete-associated landcover types. (B) Ordination of polychaete densities. Landcover type abbreviations as in Fig 1.

In regard to the differentiation of landcover types on Can1, the densities of the goniadid *Goniada japonica* and the spionid *Malacoceros indicus* were the most important variables and yet exhibited opposite effects. The density of the sabellid *Laonome albicingillum* was the third most important factor. On Can2, the densities of the nereid *Neanthes glandicincta* and the capitellid *Capitella* sp.1 played the most important roles (Fig 3A). These data reveal that each of the three landcover types was associated with a distinct polychaete species: *G. japonica* mostly occurred in the inner flat with high inundation, while *M. indicus* was primarily distributed in the mangrove area, and *N. glandicincta* was most abundant in the inner flat with intermediate inundation (Fig 3). The densities of *G. japonica* in inHF, *M. indicus* in the mangroves and *N. glandicincta* in inMF averaged 165.8, 252.1 and 712.2 individuals m^-2^, respectively (S6 Table).

### Association between landcover type and birds

The distribution of avian guilds was associated with two landcover types (Table 2). One landcover type consisted primarily of the flats, including the inner flat (inF) and the creek opening flat (croF), while the other type consisted of the mangroves (Fig 4A). In addition to this separation, further grouping indicated that the flat zone as a whole also occurred in combination with another three zones, including the intermediately vegetated sand dune (SdMV), the unvegetated sand dune (SdNV) and some locations with mangroves. Similarly, the mangrove area as a whole also included the highly vegetated sand dune (SdHV).

**Fig 4.**
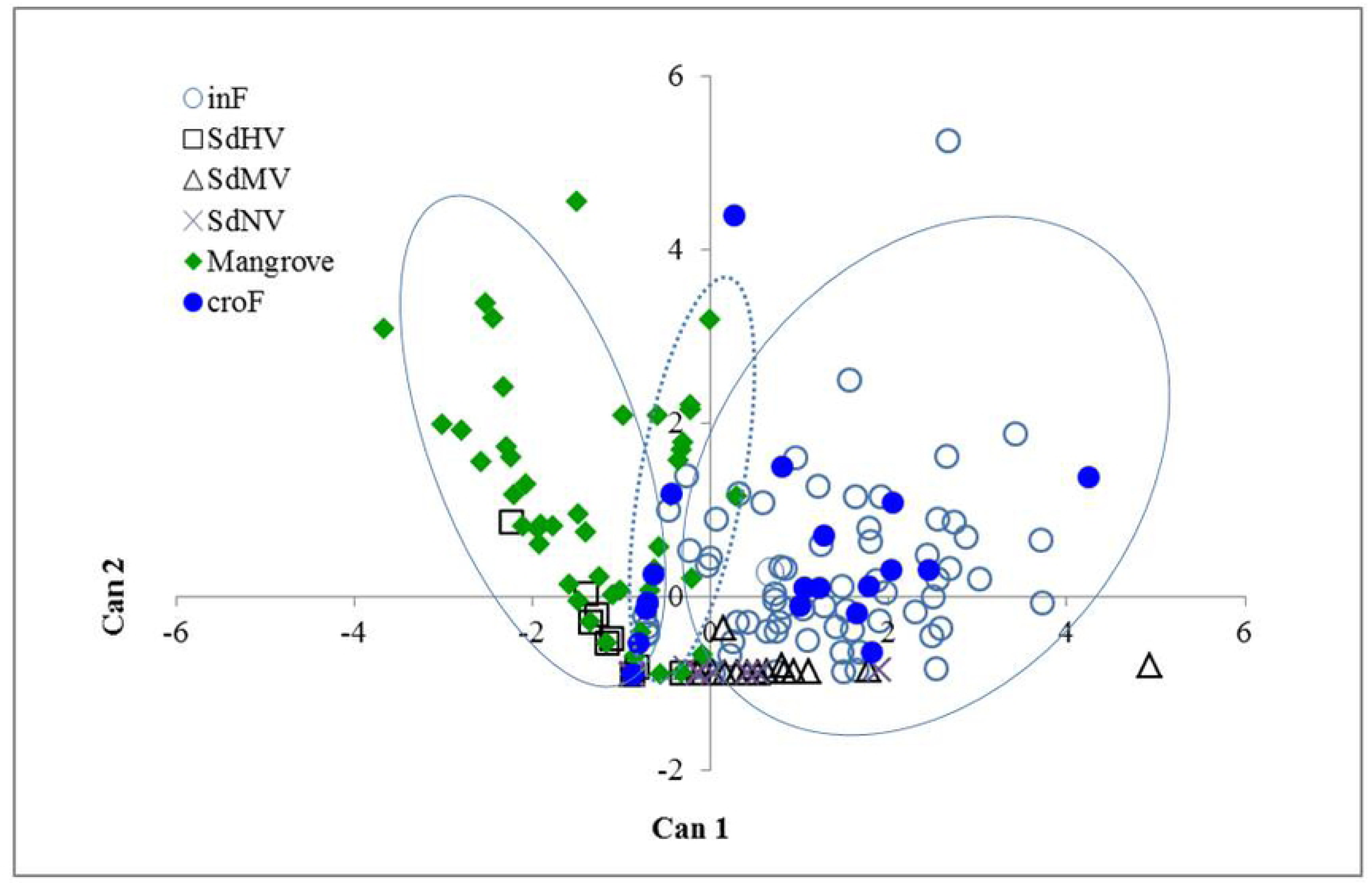
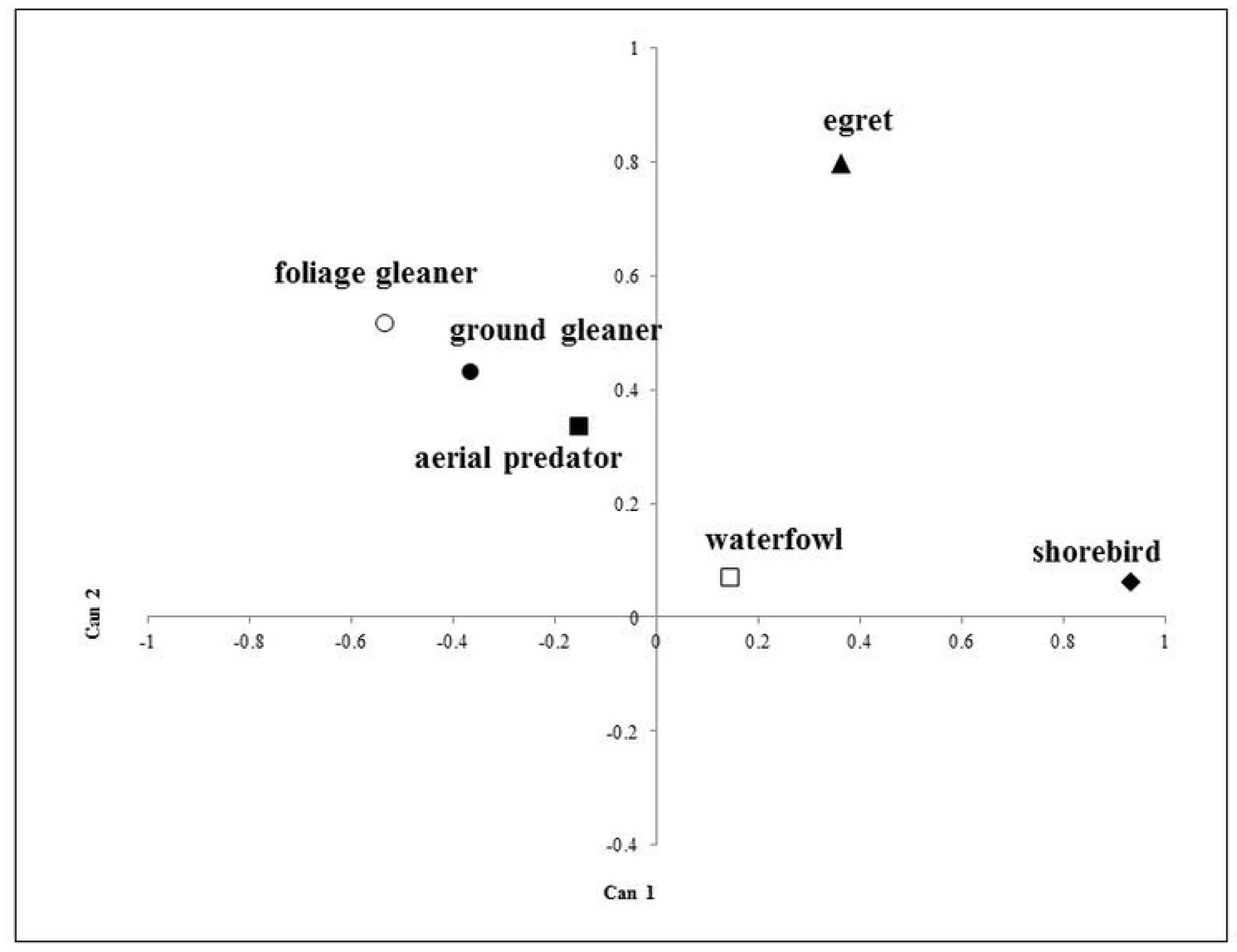
The results of canonical discriminant analysis for individuals of avian assemblage. (A) Differentiation of three bird-associated landcover types. (B) Ordination of bird counts. Landcover type abbreviations as in Fig 1. Note that some samples in mangroves overlapping with those in inner flat (inF) and creek opening flat (croF) areas correspond to egrets.

When the landcover types were separated, shorebirds had the most significant effect on Can1, while the effects of foliage gleaners were also important but had opposite effects as shorebirds. On Can2, the count of egrets had the greatest effect (Fig 4B). As a result, the two landcover types were characterized by distinguishable avian guilds, where shorebirds were dominant in the inner flat and creek opening flat while foliage and ground gleaners occurred primarily in the mangroves. Noticeably, the egrets were associated with both the tidal flat and mangroves, while the shorebirds, in one case, were exceptionally abundant in the intermediately vegetated sand dune area (Fig 4).

When the same analyses used for avian guilds were used, two landcover types were also distinguishable for avian species composition (Table 2). One landcover type consisted primarily of the flat zones, including the inner flat and the creek opening flat, while the other landcover type comprised the mangrove area. Further evaluation showed that each of these two identified landcover types occurred in combination with additional landcover types and that the types grouped into each of these two types were similar to those observed for the count attributes (Fig 5A). On Can1, the shorebirds and egrets had the most significant effect for avian species composition, while the foliage and ground gleaners were also important but had opposite effects. On Can2, the egrets, foliage gleaners and ground gleaners were the most important factors (Fig 5B).

**Fig 5.**
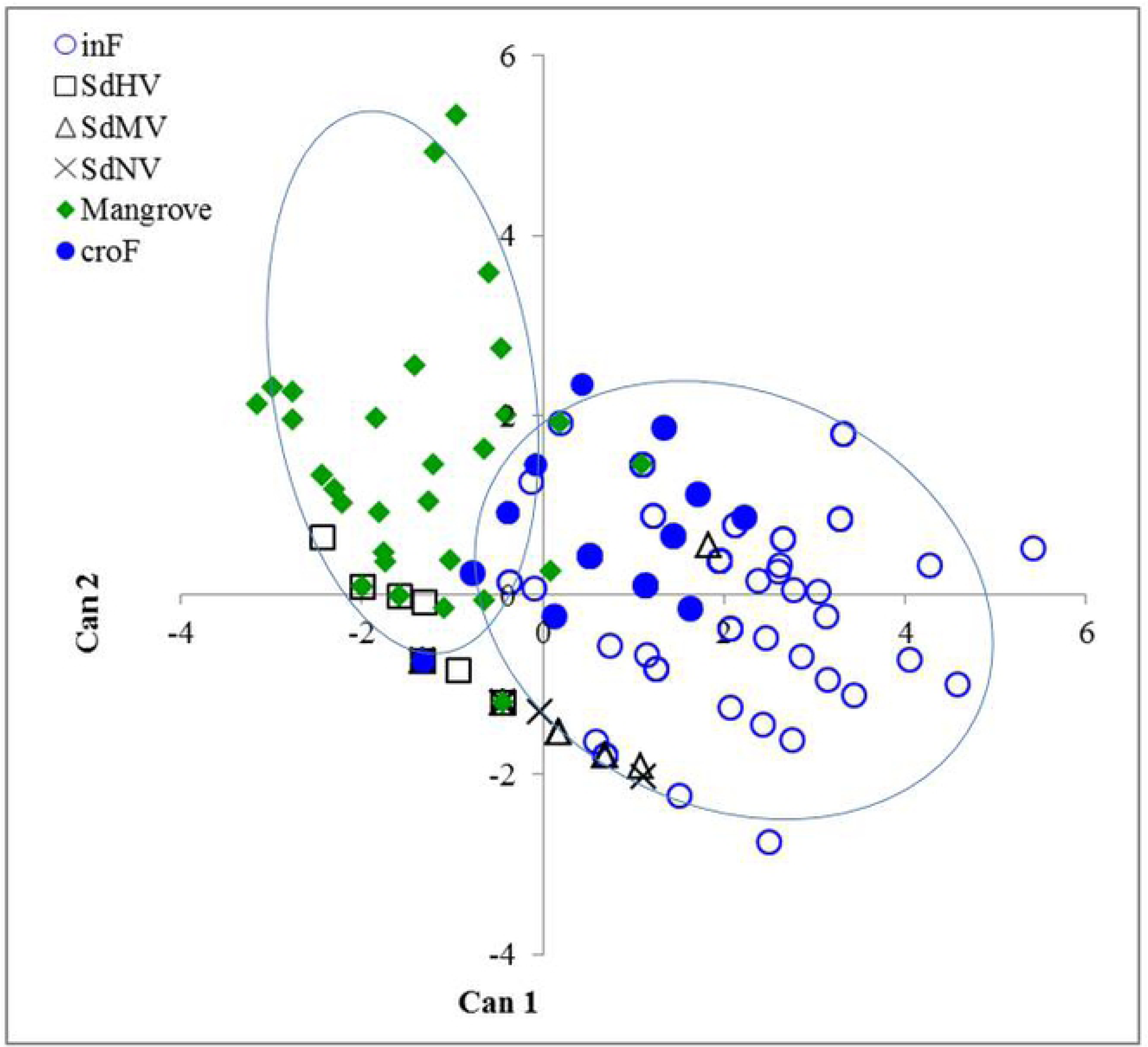
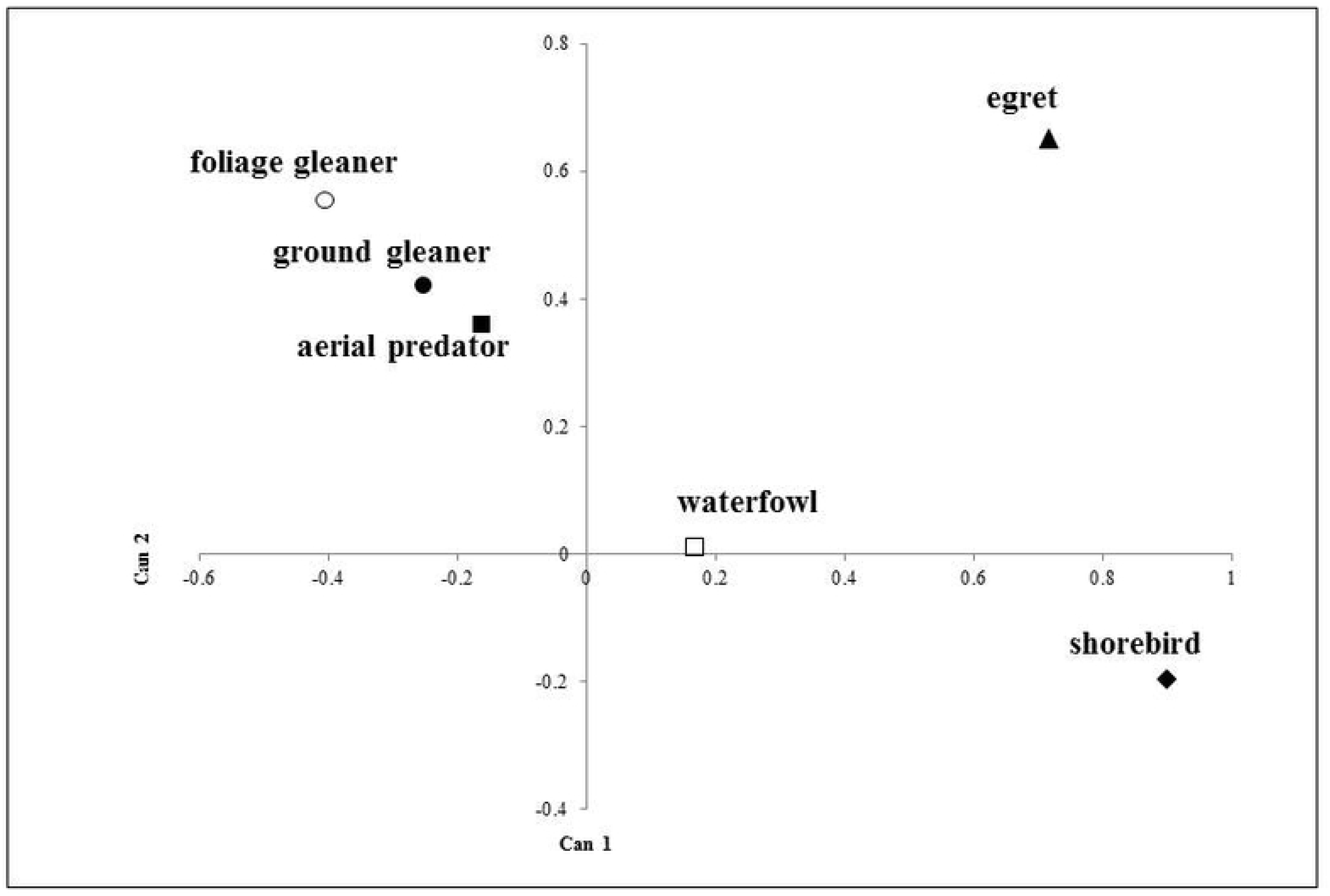
The results of canonical discriminant analysis for species richness of avian assemblage. (A) Differentiation of two bird-associated landcover types. (B) Ordination of bird species richness. Landcover type abbreviations as in Fig 1.

### Relationships among the polychaete assemblage and physical attributes

According to the CCA, six physical attributes (elevation, inundation frequency, pH, slope, flow velocity and moisture) affected the 6 polychaete species distributions (pseudo F = 4.1, p = 0.001, Table 3). The CCA ordinations revealed that *G. japonica* and *L. albicigillum* were most abundant in frequently inundated areas, while *N. glandicincta* was primarily associated with high flow velocity (Fig 6). In contrast, *P. aibuhitensis* and *M. indicus* tended to aggregate at locations with relatively high elevation and inclination. *Capitella* sp. showed a similar trend, but to a lesser extent.

**Table 3.** Summary of the results of the canonical correspondence analysis (CCA) performed on the polychaete assemblage and the redundancy analysis (RDA) performed on the bird assemblage

|  | Ordination axis |  |  |  | Total | Full model |  |  |
| --- | --- | --- | --- | --- | --- | --- | --- | --- |
|  | 1 | 2 | 3 | 4 | variance | Pseudo-F | df | p |
| *Polychaetes* |  |  |  |  |  |  |  |  |
| Eigenvalue | 0.735 | 0.212 | 0.143 | 0.047 | 3.051 | 4.1 | 5, 74 | 0.001 |
| Pseudo-canonical correlation | 0.924 | 0.572 | 0.496 | 0.314 |  |  |  |  |
| Cumulative % variance explained |  |  |  |  |  |  |  |  |
| Explained variation | 24.19 | 31.0 | 35.7 | 37.3 |  |  |  |  |
| Explained fitted variation | 63.3 | 81.5 | 93.8 | 97.8 |  |  |  |  |
| Sum of all canonical eigenvalues |  |  |  |  | 1.162 |  |  |  |
| *Birds* |  |  |  |  |  |  |  |  |
| Eigenvalue | 0.180 | 0.113 | 0.027 | 0.004 | 1.000 | 3.3 | 7, 56 | 0.001 |
| Pseudo-canonical correlation | 0.613 | 0.631 | 0.417 | 0.349 |  |  |  |  |
| Cumulative % variance explained |  |  |  |  |  |  |  |  |
| Explained variation | 18.0 | 29.3 | 32.0 | 32.3 |  |  |  |  |
| Explained fitted variation | 55.3 | 89.9 | 98.3 | 99.4 |  |  |  |  |
| Sum of all canonical eigenvalues |  |  |  |  | 0.325 |  |  |  |

**Fig 6.**
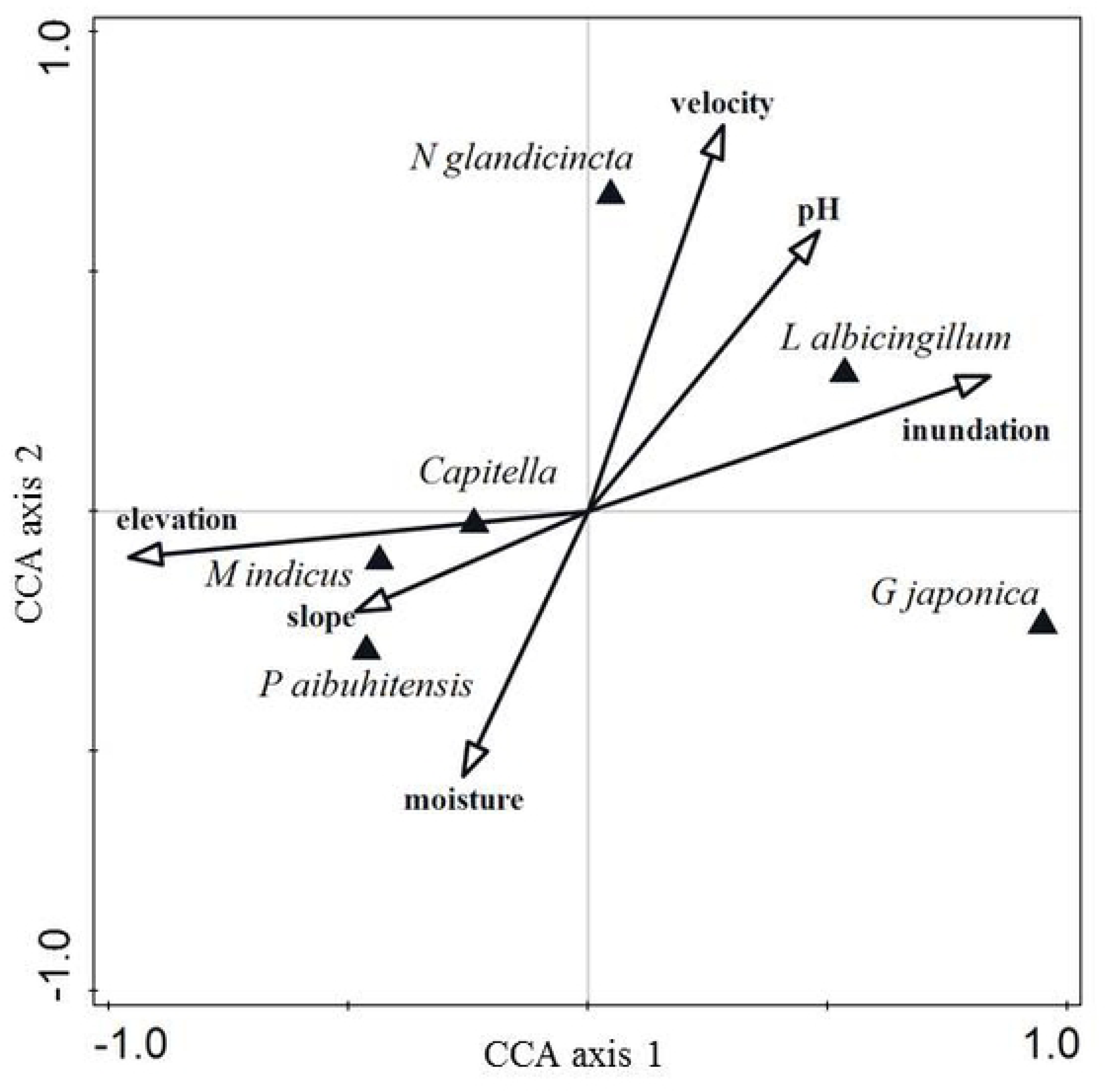
Ordination of the polychaete assemblage with physical attributes using canonical correspondence analysis. Solid triangles represent polychaete assemblage, while arrows represent physical attributes.

### Relationships among the avian assemblage and physical attributes

According to the RDA, eight physical attributes, including the surface area of the exposed open tidal flat, grain size, elevation, sorting degree, silt and clay content, inundation frequency, moisture and pH, significantly affected avian counts and species richness (pseudo F = 3.3, p = 0.001, Table 3). The RDA ordinations revealed that the shorebird counts and species richness were highest in locations with large open tidal surface areas and prolonged inundation, while the egret counts and species richness were greatest in locations with fine-grained sediment with relatively high silt and clay contents (Fig 7). In contrast, the foliage and ground gleaner counts and species richness were greatest at locations at relatively high elevation.

**Fig 7.**
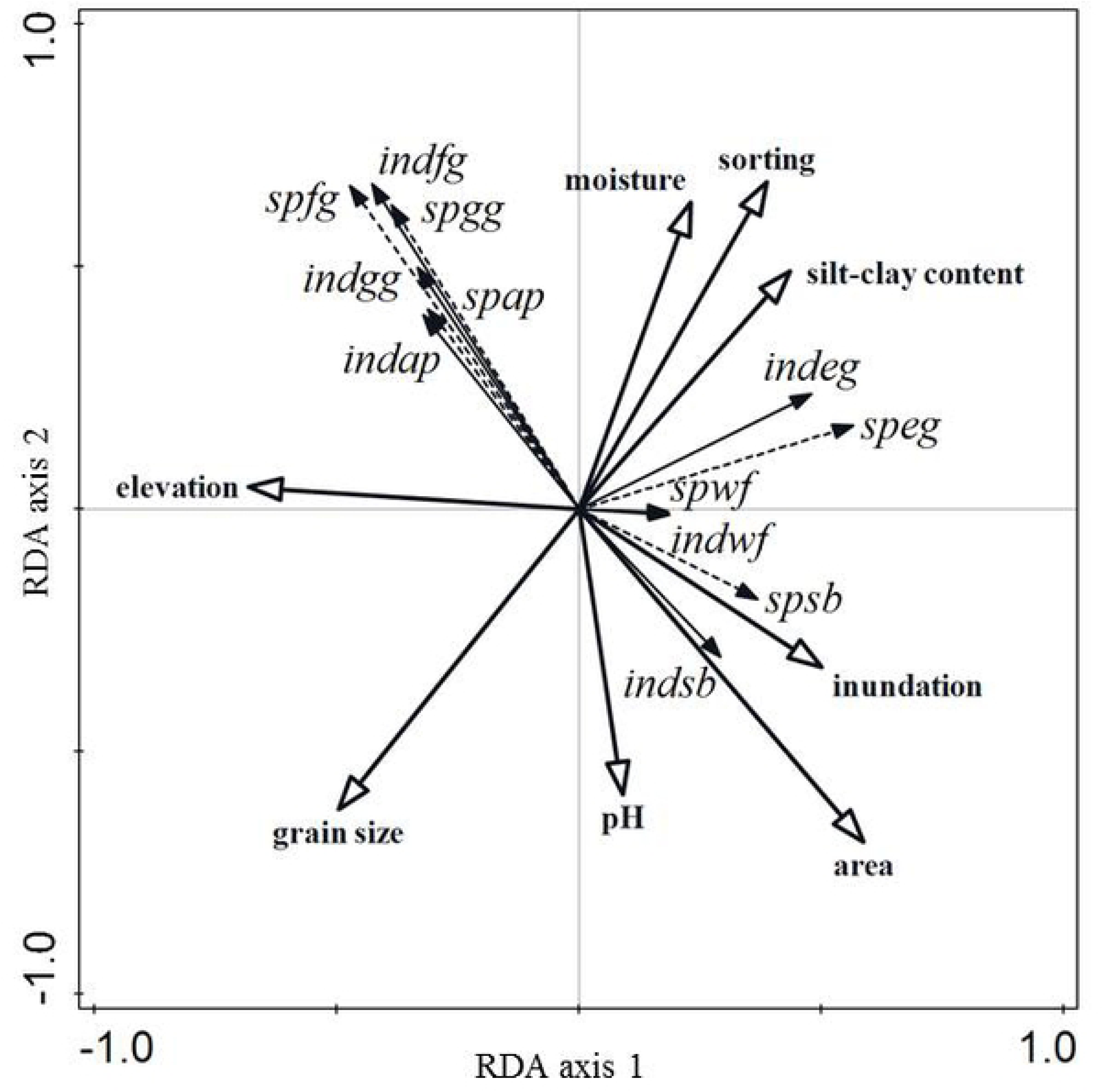
Ordination of individuals and species richness of the avian assemblage with physical attributes using redundancy analysis. Solid arrows represent avian individual while dashed arrows represent avian species richness and solid arrows with transparent heads represent physical attributes. Abbreviations are ind: individual, sp: species richness, sb: shorebird, eg; egret, wf: waterfowl, gg: ground gleaner, fg: foliage gleaner, ap: aerial predator.

## Discussion

### Variation in the composition of landcover types across the physical and biotic landscapes

Landscape structure and biotic interactions are inherently connected [14,15]. The landscape of the Wazihwei Nature Reserve, as illustrated in the present study, is similar to brocade in relation to the interplay of layers represented by the physical to polychaete and avian landscapes. The reserve encompasses six distinguishable physical landcover types, which create geomorphological, hydrological and sedimentary zonation. These physical settings serve as functional habitats for biotic assemblages, including mangroves, polychaetes and avifauna. Noticeably, the polychaetes aggregated in three fewer, but also distinguishable, landcover types, primarily the inner flat with intermediate and high inundation as well as the mangroves. This finding suggests that the polychaetes perceive the outer flat and inner low inundated flat as habitats of similar type to the mangroves. The birds explored even fewer landcover types (two), mainly the exposed open tidal flat and the mangroves. The birds appear to treat nonvegetated or intermediately vegetated sand dunes as open tidal flat, while they may perceive highly vegetated sand dunes as a habitat type similar to mangroves because both contain vegetation. The abilities of these animals to cope with the circumstances of their physical settings for their survival and foraging may highlight the differences in the physically and biologically coupled landscapes seen in the present study.

### Functions of the physical setting-polychaete assemblage-coupled landscape

Infaunal polychaetes are particle feeders and burrowers. Their feeding and burrowing activities are profoundly constrained by local flow regimes and sedimentary properties [21-23]. High silt and clay contents in sediments are associated with high organic matter (proxy of food particles) and retained water content [21, 22, 42]. Such characteristics are indicative of muddy habitats and feeble currents, which allow organic matter-laden food particles to rain down from the water column, consequently favouring deposit feeding [21, 22]. At the studied reserve, the mangrove habitat is located at relatively high elevations and has mud substrate and slow water flow, suggesting that this is a habitat favorable to deposit feeders. The spionid *M. indicus* and the capitellid *Capitella* sp. are deposit feeders [43, 44]. Their presence at high abundance in the mangrove areas of the reserve is consistent with the aforementioned expectations. The highly inundated inner flat area is also muddy, but, in comparison to the mangrove habitat, experiences prolonged submergence and faster flows. Fast flow enhances particle fluxes in the water column, which is crucial for suspension feeding [20-22]. In muddy habitats, suspension conditions can also occur when tides move over the substratum surface to resuspend food particles that were once deposited on the bottom, making food particles available to suspension feeders. The sabellid *L. albicingillum* is a suspension feeder [44, 45]. Its distribution in the highly inundated inner flat patch agrees with the expectations of feeding associated with hydrology described above.

The distribution of *G. japonica* in the inner high inundation flat of the reserve agrees with those reported in other regions where *G. japonica* is abundant at low tidal levels (northern New Zealand, [46]; western Mexico and the United States of America, Warnock N and others. unpublished data). This polychaete occurrence in a habitat with prolonged immersion perhaps correlates to its motility and carnivory. It feeds on tube-dwelling polychaetes and pericarids in the sediment [44, 47]. As lower tidal areas are subject to immersion for a relatively long duration, the sediments here become less compacted and experience unconsolidation for longer periods of time [48]. This type of habitat can benefit the foraging of *G. japonica* because the animal can move more easily and has more time to search for its prey in the sediment. We suspect that the cooccurring tube-dwelling *L. albicingillum* is its prey.

*Neanthes glandicincta* is distributed in the inner intermediate inundation flat, where the flow velocity is greatest in the reserve. This nereid is a deposit feeder and burrower [45, 49]. Fast flow together with a certain grain size (approximately 0.1 - 0.4 mm in size) can result in the transport of food particles as bedload [22, 50], thus making benthic microalgae-associated particles accessible to this deposit feeding nereid [45, 51]. In this type of flat habitat, another coinhabiting species is *Capitella* sp., which is a subsurface deposit feeder and burrower. The correlation of its distribution with fast flow might be explained in part by its specific colonization pattern at both the adult and larval stages. These worms actively recruit in the high tidal zone and soon burrow into the sediment, where the flow is faster than that in the low tidal zone [52].

### Functions of the physical setting-avifauna coupled landscape

Most shorebirds prefer foraging in unvegetated areas or those that are sparsely covered by short plants [53, 54]. The shorebirds at Wazihwei Nature Reserve were predominantly distributed in the open tidal flats. This trend reflects that shorebirds must rely on open tidal areas for foraging, a phenomenon that has been recorded extensively among most migratory shorebirds [16, 55-57]. In contrast to the large flocks of Kentish plovers (*Charadrius alexandrines*) and dunlins (*Calidris alpina*) feeding at the upper site of the inner flat with high inundation and in the creek opening flat (inFB is approximately near Station 4 in inHF and croF, see Fig 1), a very large flock of wintering Pacific golden plovers (*Pluvialis fulva*) was found in the sand dunes with intermediate vegetation (SdMV). This sand dune patch is sparsely covered by a creeping vine, the beach morning glory (*Ipomoea pes-caprae*), and a short weed plant, the black jack (*Bidens chilensis*). The small amount of vegetation in this patch results in a large beach area resembling an open sandy tidal flat, thus becoming suitable for shorebird foraging.

Egrets compose another exceptionally abundant guild in the intertidal flats. Large flocks of the little egret (*Egretta garzetta*) and the summer migrant the cattle egret (*Bubulcus ibis*) also feed in the inner flat and the creek opening flat together with the shorebirds. In addition to feeding in the open tidal flat patches, the egrets use mangrove patches as well. Little egrets, cattle egrets, black-crowned night-herons (*Nycticorax nycticorax*) and sacred ibises (*Threskiornis aethiopicus*) make nests within the mangroves (MB area, see Fig 1) during the summer from March to August. Their distributions in the reserve reveal that these four egret species share common nesting and feeding sites [27]. Furthermore, observations within a nearby mangrove reserve also within the Danshuei estuary have attributed damage to mangrove trees to droppings from and twigs trampled by nesting egrets and sacred ibises (personal communication with Pei-Fen Lee at the Institute of Ecology and Evolutionary Biology, National Taiwan University). We speculate that if nesting and roosting by these birds are intensified, the deterioration effects on the mangroves may change the landscape of this reserve.

The foliage gleaners, the light-vented bulbul (*Pycnonotus sinensis*) and the Japanese white-eye (*Zosterops japonicus*), are frugivores, nectarivores and insectivores [58-61]. Their diets consist of a variety of plants and small animals, such as fleshy fruits, flowers, dipteran insects, spiders, snails and slugs. Among the ground gleaners, the Eurasian tree sparrow (*Passer montanus*) alone accounts for 58.2% of the total ground gleaner count and is the dominant species. This species feeds mainly on wild plant seeds but shifts to insects when raising its nestlings. Dietary insects include dipterans, coleopteran adults and larvae and larval lepidopterans [62, 63]. Mangroves not only produce pollen and nectar [64] but also harbor various invertebrates, including insects. Such insects include nonbiting midges (Chironomidae), biting midges (Ceratopogonidae), flies, coleopterans and lepidopterans [65]. In addition, flowering trees, such as the native coast hibiscus (*Hibiscus tiliaceus*), are distributed on sand dunes with vegetation patches. Furthermore, seed-producing weeds, such as the common reed (*Phragmites australis*) and wedelia (*Wedelia trilobata*), cover a narrow sandy zone adjacent to the southeastern edge of the mangrove patch (zone MC, personal communication with Gwo-Wen Hwang at Hydrotech Research Institute, National Taiwan University). These vegetated patches make flowers, nectar, seeds and insects available to the foliage and ground gleaners.

### Contributions of the tidal flat and vegetation patches to the avian landscape

Most shorebirds engage in foraging at falling, low and rising tides on open mudflats [55, 66]; therefore, increases in open tidal surface area and a lengthening of time after exposure can enhance shorebird feeding success [66]. Given that the tidal flat area in the reserve (including the inner mudflat and the creek opening flat) has a large open tidal surface area and experiences the lengthening of tidal phases, the shorebirds had relatively long times and large areas over which they searched for food. As a result, the tidal flat, covering approximately 43% of the study area, represents the most vital “shorebird landscape”.

A short distance flown between a feeding and breeding site has been found in some egret species. This close spatial connection has been attributed to bird nesting success or low foraging cost [67-70]. At the present study site, egrets exhibit a spatial proximity of approximately 300 m between their feeding mudflat and nesting mangrove habitats. We suspect that this close spatial connection might attract egrets to use this reserve, particularly Little Egrets and Cattle Egrets. The combination of the tidal flat and mangrove habitats represents a unique “egret landscape”.

The mangroves in our study site consist of a single species, *Kandelia obovata*, and compose a simple vegetation structure. Joining the mangrove patch to the patches of the highly vegetated sand dune and the narrower strip adjacent to the mangrove area appears to enhance the vegetation complexity. Such a vegetation setting represents a “foliage and ground gleaner landscape”.

### Effects of mangroves on the functional landscape

In the reserve, the mangrove vegetation patch, covering approximately 34% of the whole area, is the most conspicuous landscape other than the intertidal flat. As an ecosystem foundation and engineering species, the mangroves interact with driving forces, hydrodynamics, sedimentation and topography, thus changing the landscape of the mangrove-dominated ecosystem [2, 14, 71]. Mangrove overgrowth facilitates the homogenization of landscape structure and results in lowered macrofaunal species richness in wetland ecosystems from the bottom sediment to understory and canopy layers [2, 10, 13]. Consequently, the services provided by mangrove ecosystems, particularly those that contribute to human wellbeing, are limited [9]. The question of how to enhance species richness through the diversification of habitat patches and the restriction of mangrove expansion was previously addressed in a partial mangrove removal experiment conducted in the same estuary. This field experiment demonstrated that the creation of a small patchwork of mudflats through the partial removal of mangroves dramatically increased the species richness of wintering shorebirds [2]. In cost-effective practice in Mai Po Ramsar site, WWF Hong Kong carry out mangrove management works to remove mangrove seedlings that are growing on tidal mudflat for sustaining mudflat area where benthic fauna is abundant [72].

### Recommendations for the landscape-based management

Colonization by mangroves depends on the presence of a previously existing land formation [73] while increasing siltation facilitates mangrove establishment [71, 74]. Therefore, ecological engineering approaches such as prevention of fine sediment deposition to sustain mud flats can be applied in the reserve. In addition, the understanding of why tidal flats are important needs to be delivered in public education programs. We recommend the following managerial strategies:

1. To retain unvegetated tidal muddy and sandy flats by controlling mangrove expansion can benefit the shorebird, egret and polychaete’s habitation. Mangrove trees and seedlings growing along the areas between the mangrove edge adjacent to the tidal creek should be periodically removed to create open tidal areas.
2. To enhance tidal flushing and circulation within the reserve by widening and deepening the tidal creek opening. This action prevents siltation, thus, slows down mangrove colonization.
3. To restore naturally and hydrologically driven tidal creek by meandering instead of straightening the creek. This ecological engineering approach combining with mangrove removal along the tidal creek further create more tidal flats.
4. To remove the invasive Sacred Ibis *Threskiornis aethiopicus* by attempting the methods recently delivered by the Taiwan government (https://e-info.org.tw/node/216895). The methods are such as catching eggs and chicks while adults may be removed using net traps and air gun shooting. This action helps to protect Taiwan native birds.
5. To promote environmental education and ecotour programs on the trail in the sand dune patches. Program activities include such as aesthetic experiences in the reserve landscape and the living creatures that this landscape supports.

## Conclusions

Our study in a mangrove vegetation-dominated estuarine wetland highlights the existence of a vital landscape through linkages from the physical landscape, as the foundation, to the polychaete- and bird-dependent landscapes. Among the physical landcover types, open tidal mud-sand flat is important because it supports the polychaetes and foraging shorebirds and egrets. This result reflects the effects of the physical landscape on the mediation of the polychaete and bird landscapes. Mangrove vegetation also serves as a significant habitat type but can compete with the tidal flat habitat. Therefore, effective management of the bird landscape, particularly to conserve migratory shorebirds and egrets, requires an integrated strategy that involves maintaining open tidal flats in the landscape.

## Acknowledgements

The authors are grateful to the New Taipei City Government and the Ministry of Science and Technology (MOST 103-2621-M-002 -020, MOST 106-2621-M-002 -004 -MY3), Taiwan, for funding support. The Tenth River Management Office, Water Resources Agency, Taiwan, provided the river cross-sectional bathymetry data and water level data. We also want to thank GW Hwang, CW Chang, and CY Chen, who assisted in the field.

## Supporting information captions

**S1 Table. Transformation formulae used for the physical and biotic variables measured at the Wazihwei Nature Reserve wetland in the Danshuei estuary during 2013-2015.**

**S2 Table. Counts of each species and the species composition of the avian guilds recorded in the Wazihwei wetland in the Danshuei estuary, northern Taiwan, during 2013-2015.**

**S3 Table. Means and ranges of geomorphological, hydrological and sedimentary variables measured at Wazihwei wetland in the Danshuei estuary, northern Taiwan, during 2013-2015.** N = sample size, SE = standard error.

**S4 Table. Means and ranges of geomorphological, hydrological and sedimentary variables measured in each zone of the Wazihwei Nature Reserve wetland in the Danshuei estuary, northern Taiwan, during 2013-2015.** N = sample size, SE = standard error, ranges from minimal to maximal values are shown in parentheses, inLF, inMF, inHF= inner low, intermediate and high inundation flat, respectively, Outer= outer flat.

**S5 Table. Differences in the geomorphological, hydrological and sedimentary variables between two subgroups in the inner high inundation (inHF) zone of the Wazihwei Nature Reserve wetland in the Danshuei estuary, northern Taiwan, during 2013-2015.** Examination used the Wilcoxon 2-sample test. One subgroup consists of Stations 1, 2, and 3; the other subgroup represents Station 4. N = sample size, SE = standard error.

**S6 Table. Density (mean ± SE individuals m**^**-2**^**) and density differences of 6 dominant polychaete species measured in each zone of the Wazihwei Nature Reserve wetland in the Danshuei estuary, northern Taiwan, during 2013-2014.** Density ranges from minimal to maximal values are shown in parentheses. Differences in means among zones were examined using the Kruskal-Wallis test. N = sample size, inLF, inMF, inHF= inner low, intermediate and high inundation flat, respectively, Outer= outer flat. n.s.= not significant at 0.05.

